# tDCS changes in motor excitability are specific to orientation of current flow

**DOI:** 10.1101/149633

**Authors:** Vishal Rawiji, Matteo Ciocca, André Zacharia, David Soares, Dennis Truong, Marom Bikson, John Rothwell, Sven Bestmann

**Author notes:** Corresponding author details: Vishal Rawji, UCL Institute of Neurology, London, WC1N 3BG, UK.

## Abstract

Measurements and models of current flow in the brain during transcranial Direct Current Stimulation (tDCS) indicate stimulation of regions in-between electrodes. Moreover, the cephalic cortex result in local fluctuations in current flow intensity and direction, and animal studies suggest current flow direction relative to cortical columns determines response to tDCS. Here we test this idea by measuring changes in cortico-spinal excitability by Transcranial Magnetic Stimulation Motor Evoked Potentials (TMS-MEP), following tDCS applied with electrodes aligned orthogonal (across) or parallel to M1 in the central sulcus. Current flow models predicted that the orthogonal electrode montage produces consistently oriented current across the hand region of M1 that flows along cortical columns, while the parallel electrode montage produces none-uniform current directions across the M1 cortical surface. We find that orthogonal, but not parallel, orientated tDCS modulates TMS-MEPs. We also show modulation is sensitive to the orientation of the TMS coil (PA or AP), which is through to select different afferent pathways to M1. Our results are consistent with tDCS producing directionally specific neuromodulation in brain regions in-between electrodes, but shows nuanced changes in excitability that are presumably current direction relative to column and axon pathway specific. We suggest that the direction of current flow through cortical target regions should be considered for targeting and dose-control of tDCS.

**Highlights:**

- Direction of current flow is important for tDCS after-effects.
- tDCS modulates excitability between two electrodes.
- tDCS differentially modulates PA and AP inputs into M1.

**Abbreviations:**

PA: postero-anterior
AP: antero-posterior
ML: medio-lateral
tDCS: transcranial direct current stimulation
MEP: motor evoked potential
M1: primary motor cortex
TMS: transcranial magnetic stimulation;
AP-TMS-MEPs: motor evoked potentials elicited with anterior-posterior directed TMS;
PA-TMS-MEPs: motor evoked potentials elicited with posterior-anterior directed TMS

**Funding:** This research did not receive any specific grant from funding agencies in the public, commercial, or not-for-profit sectors.

## Introduction

To date, the majority of studies in humans using transcranial direct current stimulation (tDCS) to modulate cortical function employ a bipolar electrode montage: one electrode is usually placed over the target site and the other at a distance. So, for the hand area of motor cortex (M1), a large anode is conventionally centred over the anatomical location of the “hand knob” of the precentral gyrus, with a cathode over the contralateral orbit (1). This dosing strategy, based on canonical studies by Nitsche, Paulus and colleagues on how the position of large electrodes influences population-averaged modulation of TMS-MEPs (2-4), is now widely applied for targeting diverse cortical target regions (5, 6) though rarely with consideration for nuanced dose response (7-11). Intra-cranial recordings (12, 13) and clinical imaging (14, 15), supported by current flow models (16, 17) show bipolar electrode montages produce current flow in brain regions between electrodes. Though putative brain targets between electrodes have been considered (18-20), previous tDCS studies have not systematically isolated the consequences of inter-electrode current flow.

The “inter-electrode” considerations provoke a second question. Animal studies in lissencephalic animals indicate polarity specific (anodal/cathodal) excitability changes for current directed normal to the cortical surface (2), which corresponds to current flow directed along the primary dendritic axis of cortical pyramidal neurons (21, 22). In the human cephalic cortex, such controlled stimulation cannot easily be achieved and the directions of current flow underneath an electrode are complex (23, 24). The position of primary motor cortex in the anterior wall of the central sulcus suggests that electrode montages that direct current flow perpendicular through this gyral wall (and thus predominantly along the primary dendritic axis of cortical pyramidal neurons) may optimally modulate corticospinal excitability (CSE). The second question we address here is therefore whether there are differences in the effect of tDCS on CSE when current is oriented perpendicularly across, compared with parallel to, the cortical surface at the level of the M1 hand area. To this end, we positioned tDCS electrodes 3.5 cm anterior and posterior to the hand area of M1 to direct current flow across the central sulcus (Figure 1). This means that depending on the position of the anode and cathode, current will flow through M1 in anterior-posterior (AP-tDCS) or posterior-anterior (PA-tDCS) direction, respectively. In a second condition, we positioned electrodes 3.5 cm medial and lateral to the M1 hand area to direct current flow in parallel along the cortical surface of central sulcus (Figure 1). We refer to this as medio-lateral tDCS (ML-tDCS). Motor-evoked potentials (MEPs) elicited with TMS (TMS-MEPs) were used to access CSE changes after stimulation with these two orthogonal tDCS orientations.

**Figure 1.**
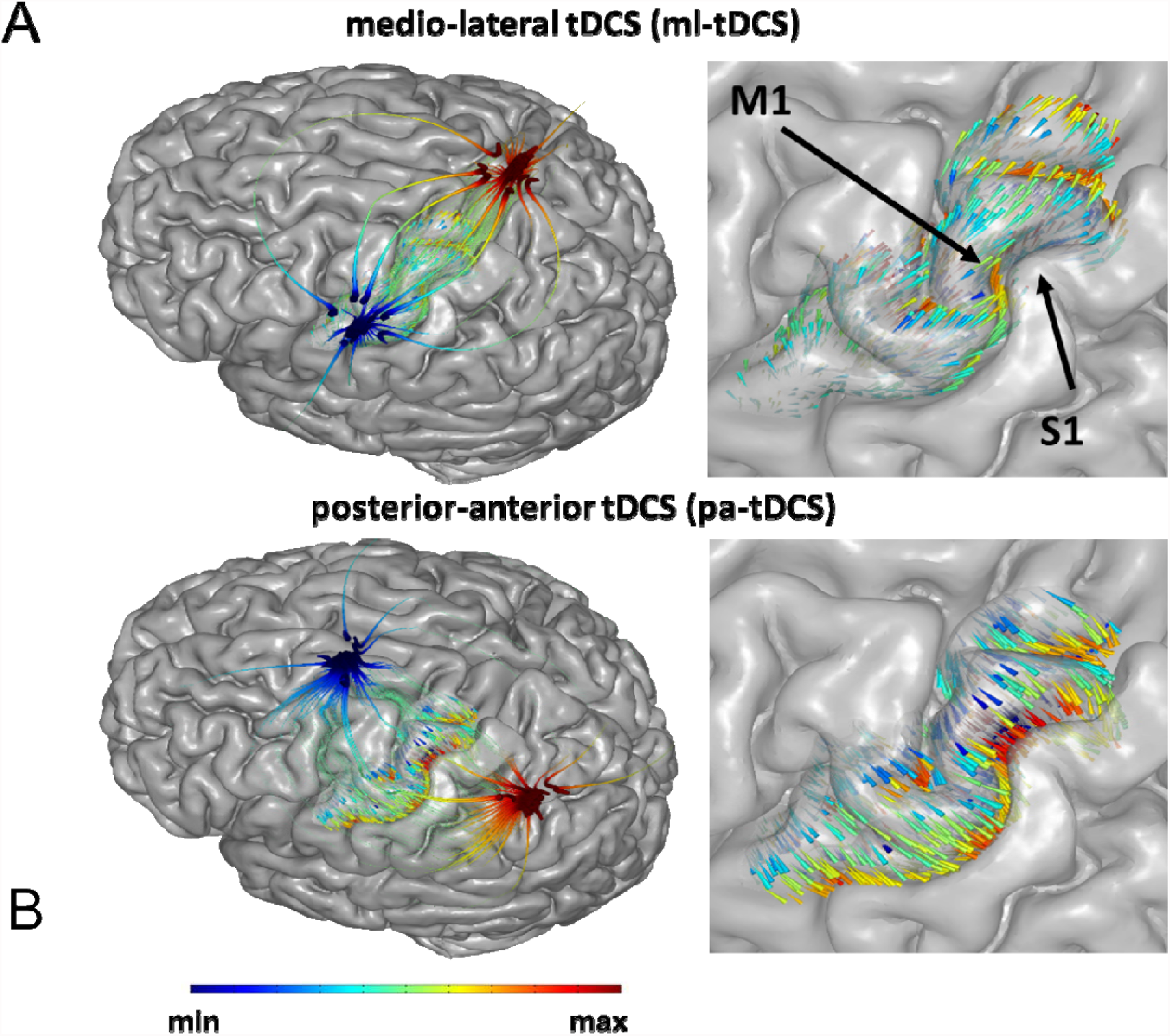
Comparison of electrical field modelling for montages directing current across and along the cortical surface. Electric field orientation on the cortex as applied by electrodes along (A) or across (B) the motor strip. Note: The streamlines and arrows have two separate colorscales. Starting outside the motor-strip, streamlines trace the direction of current density from high to low voltage (red to blue), anode to cathode. The streamlines confirm that current flows down the voltage gradient established by the electrodes and shaped by the head anatomy. On the motor strip, arrows illustrate the direction of electric field passing through the motor cortex surface. Arrow colour represents normal electric field where red is inward and blue is outward. Inward field corresponds to expected pyramidal soma depolarization and outward corresponds to expected pyramidal soma hyper-polarization (though we note that the polarization of axons and terminals must also be considered). It is important to not confuse the red/blue of voltage with the red/blue of electric field/polarization since the two are not simply related; none-the-less, this representation allows correlation of macro-scale current flow patterns set by electrode montage with gyri-scale current flow pattern which determine cellular polarization. These modelling results show how current can be directed to flow more uniformly through M1.

We also addressed a third question. The effects of TMS on motor cortex are well-known to be directional (25, 26). TMS with a monophasic pulse that induces an electric current flowing from approximately posterior to anterior across the central sulcus (perpendicular to the line of the individual’s central sulcus at that point) evokes MEPs (PA-TMS-MEPs) that have a shorter latency and lower threshold than stimulation with an anterior-posterior induced current (AP-TMS-MEPs). It is thought that this is because the two directions of stimulation activate different sets of presynaptic inputs to corticospinal neurones (21). Indeed, brain slice studies of tDCS established modulation varies across afferent axonal pathways or varied orientation (24, 27). We therefore hypothesised that any effects of tDCS across M1 might also be directionally selective, and that they would interact in different ways with the direction of TMS used for eliciting MEPs. Specifically, using tDCS to direct current perpendicular to the M1 hand area, we expected that stimulation with a posterior cathode and anterior cathode (PA-tDCS) would influence MEPs evoked by PA and AP TMS in a different way to tDCS applied with an anterior anode and posterior cathode (AP-tDCS).

## Methods

### Participants

22 healthy volunteers (17 male, 21 right handed) aged 21-44 (mean age 28.95, SD 6.14) participated in this experiment. The study was approved by the UCL Ethics Committee and none had contraindications to TMS or tDCS as assessed by a TMS/tDCS screening questionnaire.

### Current flow modelling

Finite Element Method (FEM) models of tDCS were generated to predict electric field (E-field) orientation along the motor cortex. High resolution T1 and T2 weighted MRI scans (GRE sequence, TR = 1900ms, TE =2.2 ms and SPACE sequence, TR = 3200 ms, TE = 402 ms respectively) were previously collected and segmented using a combination of automated and manual segmentation techniques (28). Automated segmentation algorithms derived from Unified Segmentation in SPM8 (29, 30) were combined with updated tissue probability maps and morphological filters (smoothing, dilation, erosion) specifically developed for current flow modeling (31). Additional image masks (fat, electrodes, gels) and regions of interest (M1) were segmented using manual and semi-manual tools (Simpleware, Synopsys) to remove aliasing artifacts, incorporate gyri-precise detail, and position stimulation electrodes along or across the hand regions of the motor cortex, as identified using anatomical landmarks (32, 33). Adaptive tetrahedral meshes were generated using a voxel-based algorithm (Simpleware, Synopsys) with multiple domains corresponding to different material conductivies verified by intra-cranial recording (in S/m: Scalp 0.465, fat 0.025, skull 0.01, CSF 1.65, Grey matter 0.276, White matter 0.126, Air 1e-15, electrode 5.99e7, gel 1.4) (12, 34, 35). Meshes were imported into a FEM solver (COMSOL Multiphysics) where the Laplace equation for electrostatics (∇ · (σ∇V) = 0) was solved as the field equation given a normal current density boundary condition on the anode equivalent to 1mA, ground boundary condition on the cathode, and insulation on all other external boundaries. Results were scaled linearly to match experimental conditions when necessary. E-field orientation was visualized with surface arrows seeded evenly along M1. Arrow colors corresponded to E-field normal (n·E) to the cortical surface. By convention, positive normal E-field represented inward “anodal” E-field (red) while negative normal E-field represented outward “cathodal” E-field (blue). Streamlines representing current flow through M1 were generated by seeding 100 points randomly along the surface of M1 and the gel-skin contact. Current density was then traced throughout the model. Line thickness was a logarithmic function of current density magnitude and colorized to Voltage (red, anode; blue, cathode).

### EMG Recordings

Throughout the experiment, subjects were seated comfortably in a non-reclining chair, with their right hand rested on a cushion. Electromyographic (EMG) activity was recorded from the right first dorsal interosseous (FDI) muscle using Ag/AgCl cup electrodes arranged in a belly-tendon montage. The raw signals were amplified and a bandpass filter was also applied. (20Hz to 2kHz (Digitimer, Welwyn Garden City, UK)) Signals were digitised at 5kHz (CED Power 1401; Cambridge Electronic Design, Cambridge, United Kingdom) and data were stored on a computer for offline analysis (Signal Version 5.10, Cambridge Electronic Design, UK was used).

### Transcranial Magnetic Stimulation

Single pulse TMS was employed using a Magstim 200^2^ stimulator (The Magstim Co. Ltd) connected via a figure-of-eight coil with an internal wing diameter of 7cm. The hotspot was identified as the area on the scalp where the largest and most stable MEPs could be obtained for the right FDI muscle, using a given suprathreshold intensity. The coil was held approximately perpendicular to the presumed central sulcus and held tangentially to the skull with the coil handle pointing backwards for postero-anterior (PA) stimulation and handle pointing forwards for antero-posterior (AP) stimulation. A coloured pencil was used to draw the boundaries around the coil so that it could be accurately positioned to the hotspot for further recordings, for PA and AP coil orientations.

### Transcranial Direct Current Stimulation

Transcranial direct current stimulation (tDCS; Starstim, Barcelona; 1 mA) was applied via 3.14 cm^2^ Ag/AgCl gelled electrodes yielding an average electrode current density of 0.318 mA/cm^2^. The stimulation was applied for a total of 10 min, ramped up and down for five seconds at the beginning and end of the stimulation. Participants were asked to stay awake and at rest during the stimulation. Sham stimulation involved ramping up then down both at the start and end of the 10 min period, with zero stimulation for the remaining time.

TDCS electrodes were positioned 3.5cm anterior and posterior to the TMS hotspot along the orientation of the TMS coil; two further electrodes were placed 3.5cm medial and lateral to the hotspot perpendicular to the coil orientation (i.e. along the length of the central sulcus). We note that this convention is with regards to the approximate orientation of the central sulcus, which is generally oriented at about 45 degrees with respect to the midline. For simplicity, we will assume that this corresponds to the anterior-posterior and medio-lateral orientation of the brain.

Stimulation was set up remotely on a computer and delivered via a Bluetooth receiver connected to the electrodes. We refer to stimulation with a posterior anode and anterior cathode as PA-tDCS, to indicate the direction of the electric field; AP-tDCS refers to stimulation with an anterior anode and posterior cathode. We refer to stimulation with a medial anode and lateral cathode (directing current flow approximately in parallel to the central sulcus) as ML-tDCS (see Figure 2).

**Figure 2.**
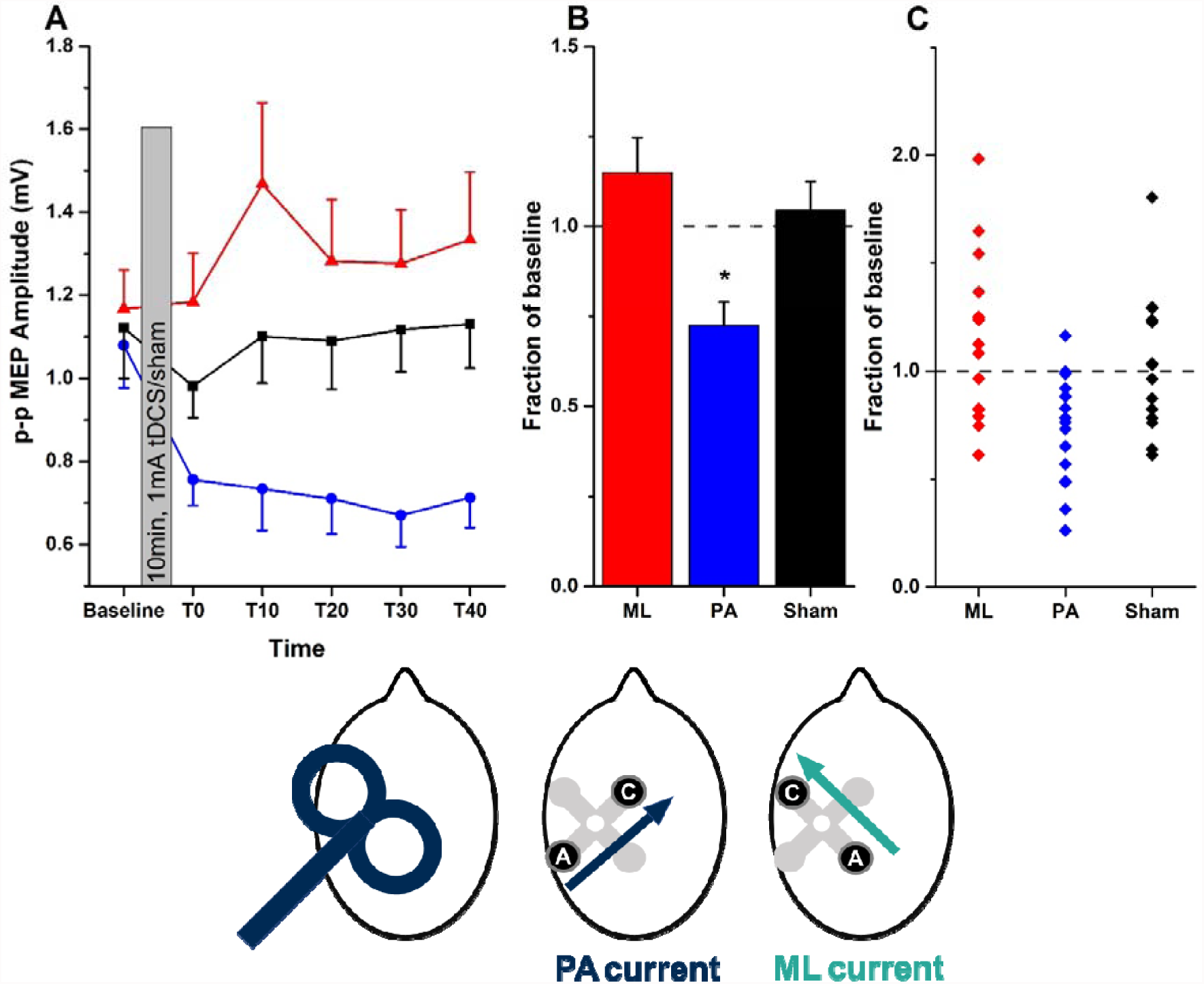
Effect of PA- and ML-tDCS on the amplitude of MEPs evoked by PA-TMS-MEPs. **A**, mean (±SEM) MEP amplitudes at baseline and every 10 minutes between T0 and T40 (red triangles = ML-tDCS, black squares = Sham, blue circles = PA-tDCS). **B,** mean (±SEM) post-stimulation effect (averaged from T0 to T40) expressed as a fraction of the baseline value in each group (ML, red; PA, blue; sham, black). Asterisks represent paired t-test significant differences (p<0.05 with Bonferroni’s multiple correction) when compared to sham. **C,** individual data points contributing to the mean values plotted in B. The head diagrams represent coil orientation and electrode configuration used: PA-TMS-MEPs, PA-tDCS and ML-tDCS.

### Experimental Parameters

Resting motor threshold (RMT) was defined as the lowest TMS stimulus intensity to evoke a response of 50 μV in 5 out of 10 trials in the relaxed FDI using the optimal PA orientation.

Twenty MEPs were collected before and after tDCS, with post tDCS MEPs collected every ten minutes from T0-T40. The TMS stimulus intensity was set at the intensity required to evoke a response of 1 mV peak-to-peak amplitude (SI1mV), and this intensity was kept constant throughout the entire experiment. The mean amplitude of these MEPs was calculated for each time point for each subject.

### Experiment 1

In experiment 1, we investigated whether corticospinal excitability could be modulated with an electrode montage for which the region of interest (M1 hand region) was positioned between our stimulating electrodes. This was accomplished using two different stimulating montages, PA and ML tDCS (as described above), to investigate whether direction of current flow across M1 could differentially modulate responses. To this end, fifteen people participated in a crossover study, which consisted of three randomised sessions (ML-tDCS, PA-tDCS and sham), each separated by at least five days. MEPs were assessed before and after tDCS using PA TMS.

### Experiment 2

Following the interesting results from experiment 1, we next asked whether the observed effect of PA-tDCS on corticospinal excitability might be explained by specific modulation of posterior-to-anterior or anterior-to-posterior inputs into M1, as probed by exploiting the known directional sensitivity of TMS over M1. Fourteen people participated in this experiment. PA-tDCS was applied and the effects on PA-TMS-MEPs and PA-TMS-MEPs were assessed. For each TMS stimulus direction at each time point, 20 MEPs were acquired.

### Experiment 3

To complete the permutations, we finally assessed the effect of AP-tDCS on both PA and AP TMS pulses in the same group as in experiment 2.

## Data Analyses

The amplitude of each single MEP at each time point was measured and averaged in each individual. Data from each individual was then averaged into a grand mean and entered into a two-way repeated measures analyses of variance (rmANOVA) with main factors “STIMULATION” (in experiment 1: ML-tDCS, PA-tDCS and Sham) or “COIL DIRECTION” (PA and AP TMS coil orientations in experiments 2 and 3) and “TIME” (Baseline, T0, T10, T20, T30 and T40 for all experiments). Absolute MEP values were used in each statistical test. In cases where there was a significant “STIMULATION” x “TIME” OR “COIL DIRECTION x TIME” interaction, analysis showed no overall effect of TIME from T0-T40 (i.e. post-tDCS). Therefore we calculated the mean post-tDCS effect and expressed this as a fraction of the baseline for post hoc testing. To examine test-retest reliability between individuals having repeated PA-tDCS sessions, we used a (2,k) intraclass correlation coefficient.

## Results

### Modelling Current Flow

The electrode montages we used have not been explored in detail previously. Adapting models with parameters previously validated by intra-cranial recording (13), we calculated the expected electric field distribution in the central area of the cerebral cortex using the electrodes and stimulation sites in the present study. Figure 1 shows the predictions for ML-tDCS (top) and PA-tDCS (bottom). As reported by others (17, 31, 36, 37), the modelling shows that with bipolar montages substantial current flow (field intensities) are produced between the two electrodes. There is a notable and clear difference between the two electrode montages: whereas ML-tDCS does not produce any uniformly directed electrical fields through the surface of motor cortex, PA-tDCS leads to relatively uniform inward and outward electrical fields, which are perpendicular relative to the cortical surface of M1. For AP-tDCS the current directions reverse (not shown). Based on data from animal models (22), inward and outward electric fields would correspond to preferential pyramidal soma depolarization and hyper-polarization, respectively. For ML-tDCS, the direction of the electric field is predominantly parallel long the cortical surface in central sulcus, which would suggest no dominant pyramidal soma polarization polarity. A following prediction would be that electrode montages that lead to relatively uniform electrical fields directed perpendicular to the cortical surface in M1 should be more efficient in modulating CSE.

### Physiological Measurements

No significant differences were found between TMS thresholds or amplitudes of the test MEP across sessions. As expected AP TMS thresholds were higher than those for PA stimulation (see Table 1).

### PA-versus ML-tDCS: effects on PA-TMS-MEPs

15 individuals were given 10 min of 1 mA tDCS using each of three separate tDCS montages tested in separate sessions at least one week apart: PA-tDCS, ML-tDCS or sham-tDCS. MEPs were evaluated at rest before and up to 40 min after tDCS (Figure 2A). PA-tDCS decreased the amplitude of MEPs whereas there was no effect of either sham or ML-tDCS. This was confirmed in a two-way repeated measures ANOVA with STIMULATION (PA, ML, sham) and TIME as main factors which showed a significant STIMULATION x TIME interaction (F (10, 140) = 1.991; p = 0.039, η^2^*=* 0.125) indicating that MEPs were affected differently by each type of tDCS. In order to understand the source of the interaction, the mean post-tDCS effect has been expressed as a fraction of the baseline in Figure 2B. A one way ANOVA showed a significant effect of type of STIMULATION (F (2, 28) = 7.134; p = 0.002, η^2^ = 0.311), with posthoc (Bonferroni corrected) paired t-tests and effect sizes showing a significant difference between the effect of PA-tDCS and sham (t = −3.279; p = 0.005; d = −1.110) and PA v. ML-tDCS (t = −3.196; p = 0.006) but not between sham and ML-tDCS (t = 0.760; p = 0.46; d = 0.302).

### PA-tDCS: contrasting effects on PA- and AP-TMS-MEPs

We next examined in 14 participants whether PA-tDCS had different effects on the MEPs evoked by TMS directed in AP or PA fashion (ie. the coil handle pointing forward or backward). Baseline MEPs to each direction of TMS had the same amplitude (AP-TMS-MEPs: 0.983mV ± 0.201 vs AP-TMS-MEPs: 1.042mV ± 0.226), although the absolute intensity required for AP-TMS-MEPs (49.1 ± 2.6%) was higher than for AP-TMS-MEPs (62.9 ± 2.8%). Collapsing all the post-tDCS MEPs and expressing them as a fraction of the baseline (Figure 3A) revealed a significant effect of PA-tDCS on AP-TMS-MEPs (t = −2.73; p = 0.017, d = −0.980) but no effect on AP-TMS-MEPs. There was a significant difference in the effect on PA-TMS-MEPs vs AP-TMS-MEPs (t= −2.565; p = 0.024, d = −0.946).

**Figure 3.**
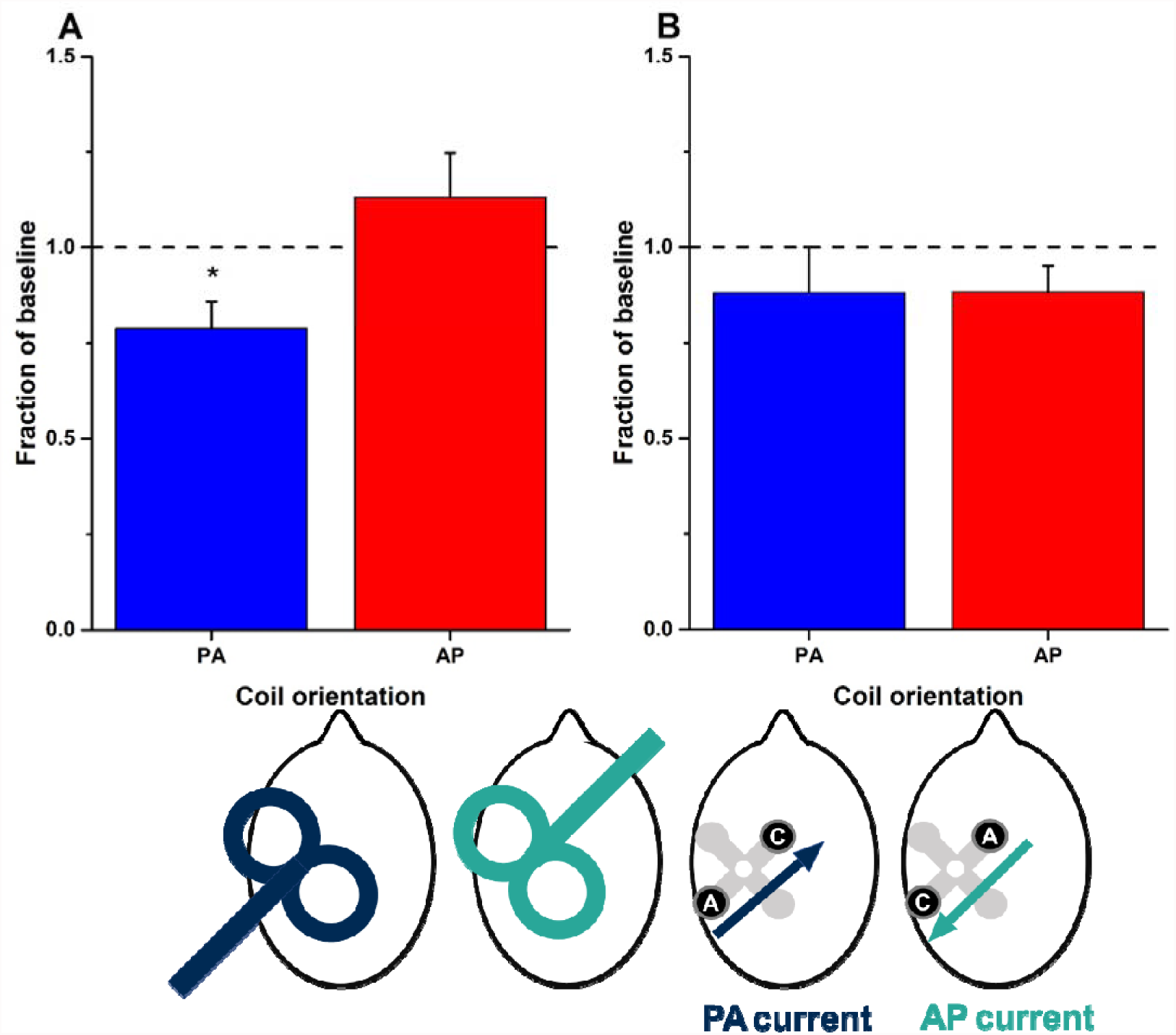
Effect of PA-tDCS (A) and AP-tDCS (B) on the amplitude of MEPs evoked by PA- and AP-TMS-MEPs. Graphs plot overall mean (±SEM) post-stimulation effects (averaged from T0 to T40 and expressed as a fraction of baseline values) for PA- (blue bars) and AP-TMS-MEPs (red bars). Asterisks represent significant differences between the two coil orientations (p<0.05 with Bonferroni’s multiple correction). The head diagrams represent coil orientation and electrode configuration used: PA- and AP-TMS-MEPs, and PA- and AP-tDCS.

### AP-tDCS: contrasting effects on PA- and AP-TMS-MEPs

Given the equivocal effects of PA-tDCS on AP-TMS-MEPs (experiment 2), we finally tested whether more consistent effects might be observed on AP-TMS-MEPs when AP-tDCS was employed. In the same participants as in experiment 2 we compared the effects of AP-tDCS on PA-TMS-MEPs and AP-TMS-MEPs (Figure 3B). Collapsing all the post-tDCS MEPs and expressing them as a proportion of the baseline, post hoc paired t-tests revealed that there was no difference in the effect of AP-tDCS on MEPs evoked by the two coil orientations (t = −0.112; p = 0.913, d = 0.035) nor were the MEPs to either type of stimulation changed in size after AP-tDCS (Figure 3B).

### Variability in responses to PA-tDCS

In total, 22 different individuals were examined for the effects of PA-tDCS on PA-TMS-MEPs. The results for all of them are plotted in Figure 4A to illustrate the variability of the effect. Averaging over all post-tDCS time points gives a mean reduction of the MEP to 74.3 ± 5.1% of baseline values.

**Figure 4.**
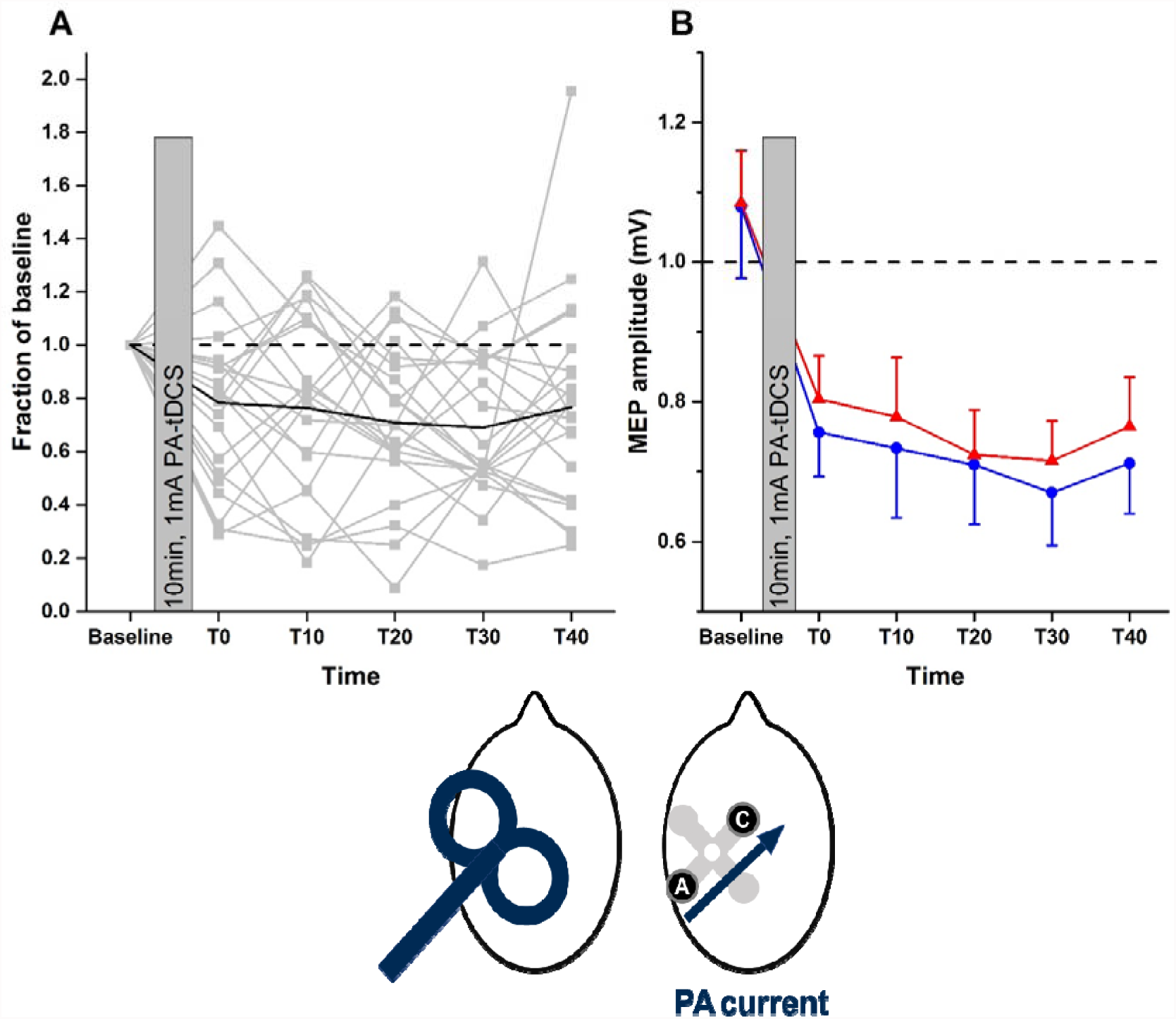
A, inter-individual variation in the effect of PA-tDCS; B, repeatability of group mean response to PA-tDCS on the amplitude of MEPs evoked by PA-TMS-MEPs. **A**, In all 22 individuals, MEPs evoked at each post-TDCS time point have been expressed as a fraction of baseline. The solid black line represents the average response from all individuals. **B**, mean (±SEM) amplitudes at baseline and every 10 minutes, post-tDCS in a group of 15 participants who were tested on 2 separate occasions (red and blue symbols and lines). The head diagrams represent coil orientation and electrode configuration used: PA-TMS-MEPs and PA-tDCS.

In addition, 15 of these participants were tested on two separate occasions. The mean data from session 1 and 2 are shown in Figure 4B. A two-way ANOVA showed no effect of SESSION (p=0.2, F (1, 14) = 1.808; p = 0.2, η^2^ = 0.028) and no SESSION*TIME interaction (F (5, 70) = 0.555; p = 0.70, η^2^ *=* 0.223) indicating no significant differences between the two sessions. There was however, a main effect of TIME (F (5, 70) =7.328; p = 0.017, η^2^ = 0.630). The interclass correlation coefficient for the mean percentage reduction in MEP was 0.68, which is generally classified as moderate-strong reproducibility.

## Discussion

Selecting the scalp positon of electrodes relative to a brain target is a paramount consideration in tDCS (5) – the common bipolar montage places one electrode over the target and the second electrode at some distance. Motivated by previous studies (see below) and specific modelling predictions (Figure 1), the present study shows that bipolar tDCS produces quantifiable changes in the excitability of primary motor cortex when it is located between the positions of the two stimulation electrodes, as opposed placing one electrode over M1. For the hand area, the effects on corticospinal excitability depends also on the direction of the electric field: in line with the predictions from current modelling, a montage orthogonal (perpendicular) to the gyrus generated consistently directed electric fields as compared with a montage along (parallel) to the gyrus. This was reflected in the after-effects on MEPs: they were suppressed after delivering perpendicular current but there was no effect after parallel current. Finally, the direction in which tDCS was applied across the sulcus (i.e. anode anterior or posterior) interacted differentially with the direction of TMS pulses: PA-tDCS affected PA-TMS-MEPs, but had no effect on AP-TMS-MEPs. AP-tDCS had no significant effect on either direction of TMS.

### Stimulation between two electrodes

Previous modelling studies have predicted bipolar tDCS will produce cortical electric fields between the two stimulation sites which may be as large or larger than those immediately under the electrodes (17, 38, 39) – predictions recently validated (13, 40, 41). Moreover, our own modelling presented here suggests that perpendicular current flow through the sulcus produces more uniformly directed current at the target site, compared to montages that direct current along the sulcus. This leads to the prediction that tDCS of M1 hand area will have different effects when current is passed between electrodes posterior and anterior to the axis of the central sulcus, compared with electrodes placed medial and lateral. These predictions were indeed borne out by the present results: PA-tDCS leads to aftereffects on corticospinal excitability whereas ML-tDCS has no effect. This also indicates that not just the strength of current but also its direction with regards to the cortical surface play an important role in mediating changes in corticospinal excitability.

Although it is possible that the posterior anode changes activity in parietal cortex and this secondarily leads to changes in M1, previous work suggests that electrodes positioned more posteriorly do not effectively modulate corticospinal excitability (36). We conclude that stimulation of sites between two scalp electrodes occurs with bipolar tDCS, as dictated by the physics of current flow, which has important implications for studies seeking to target a specific brain region. Moreover, our results indicate that controlling for the direction of current flow through a target region may help to improve the efficacy of tDCS. For example, it is conceivable that in many studies a mix of PA and ML currents will occur between subjects. If one adds to this the notion that the intensity of stimulation will vary greatly across subjects when controlling stimulator output rather than effectively applied current inside the brain (42, 43), we have the situation of large variability in applied current intensity and direction at the presumed target site. Indeed, given our results, in which ML-tDCS produced no reliable effects on CSE, such a mix of current flow direction may contribute to reports of inter-subject variability in physiological and behavioural stimulation outcomes (44-47). At least for motor cortex, control of current flow direction could be easily achieved based on the optimal orientation and position of TMS for eliciting motor-evoked potentials.

### Directionality of tDCS directed across central sulcus

“Anodal Stimulation” with a large electrode placed directly over M1 can increase cortical excitability. This is usually explained in the following way. Anodal stimulation produces in inward current flow (18, 37), though not exclusively depending on cortical folding (24). Cortical pyramidal neurons, including in M1, are aligned perpendicular to the surface of the cortex, such that an inward current flow hyperpolarises their dendrites and depolarises the cell body (22). Neurophysiological studies in animal indicate that the net effect of this is an increase the excitability of the neuron (48, 49) including to synaptic inputs (24), favouring build-up of an LTP-like effect over the 10 min of tDCS (2, 50). This is thought to result in larger MEPs when the same inputs are activated using TMS.

The current flow modelling work presented here shows that similarly directed tDCS produces directional current which, depending on the polarity, enters and exits from the posterior and anterior banks of the precentral gyrus. For PA-tDCS, inward flow on the anterior bank of the central sulcus should polarise the pyramidal neurones in the sulcal wall (which are oriented parallel to the surface of the brain) in the same way as direct “anodal stimulation” over M1 is presumed to operate (though we note this inference stems from animal studies with well-controlled current flow). If so, then we might expect that PA-tDCS has the same effect as conventional anodal tDCS. In fact, the opposite was observed here: 10 min of 1 mA PA-tDCS suppresses MEPs, whereas “anodal stimulation” directly over M1 enhances MEPs, despite this effect also being variable (44).

The different excitation by “anodal tDCS” applied with conventional or focal 4x1 HD electrodes (51), and inhibition by PA-tDCS may be explained by difference in which neuronal elements are modulated. In contrast to the above proposed action on cortical neurons in the gyri-wall by PA-tDCS, “anodal tDCS” may modulate TMS responses by polarisation of cortical neurons specifically in gyral crowns, where inward direct current is indeed more likely (24). PA-tDCS will produce current at the gyri-crown parallel to the cortical surface, which is orthogonal to cortical neurons but aligned with cortico-cortical axons afferents. This leads to an alternative hypothesis where PA-tDCS modulates TMS response by polarisation of afferent axons in the gyri crown, which are sensitive to direct current. Animal neurophysiology suggests direct current orientation toward the activated axon terminal (PA in this case) will indeed decrease excitability (24, 27).

A range of further alternative explanations can be proposed, that to varying extents explain the lack of the expected modulation of directional tDCS on changes in PA-TMS-MEPs or AP-TMS-MEPs. Even using HD electrodes, tDCS is not focal so net changes motor excitability may reflect actions on other cortical regions (24, 52). A none-trivial dependence on tDCS polarity was already known (8) and the non-linear properties of neurons (53) and networks can produce preferential responses to one tDCS polarity (54).

In general, the nuance in neuromodulation identified here derives from details of cortical folding and cellular morphology. Given that AP-TMS-MEPs and PA-TMS-MEPs activate different inputs to corticospinal output neurones (21) it is not unforeseen that AP-TMS-MEPs were unaffected by PA-tDCS. The dependence on idiosyncratic anatomy (and gradation in coil positioning) may lead to inter-individual variability that masks population effects for many conditions, and produce individual variability.

### Variability

We examined the after-effects of PA-tDCS on PA-TMS-MEPs in 22 individuals and found that in 15 of them tDCS reduced corticospinal excitability by 10% or more. Furthermore, the ICC for repeated assessments within an individual was 0.68. Both of these figures are higher than previously reported for standard montage tDCS (55). It would need a larger study to power this comparison adequately, but it could be that by aligning the tDCS to the individual best direction for PA-TMS-MEPs we achieved an increased uniformity of electric field normal to the surface of M1 compared with a single central anode.

### Conclusion

We provide strong support for the notion that bipolar tDCS has effects on cortex between the primary sites of stimulation, as predicted by models of current flow in the brain. Furthermore, these effects can be directionally-dependent. These factors may be important when interpreting (and comparing) results from conventional tDCS. More generally, our results indicate how current flow models can guide electrode placement and motivate experimental questions concerning the key factors for optimizing tES.

